# Visual input drives diverse ER calcium signals in neurons *in vivo*

**DOI:** 10.64898/2025.12.18.695086

**Authors:** Katherine C. Delgado, Kwass Wass, Lucie Lefebvre, Ketan R. Kotla, Erin L. Barnhart

## Abstract

The endoplasmic reticulum (ER) has long been thought to shape calcium signals in neurons, but stimulus-driven ER calcium fluctuations have not been directly measured *in vivo*. To measure neuronal ER calcium signals *in vivo*, we paired visual stimulus presentation with two photon imaging of ER and cytosolic calcium reporters in four different cell types in the *Drosophila* visual system. We found that visual input elicits diverse ER calcium signals, with the ER acting as a calcium sink or source depending on the cell type, subcellular compartment (dendrite versus axon), and type of visual stimulus. ER calcium signals were not simply a reflection of cytosolic signals, indicating that the ER, rather than acting as a passive calcium buffer, actively processes calcium signals in neurons in a context-specific fashion. Thus, ER-based signal processing may contribute to functional diversity across neuronal cell types, thereby enhancing the computational capacity of neural circuits.

## INTRODUCTION

“The neuron” is not a monolith. Neuronal cell types are morphologically, molecularly, and functionally diverse^1–6^, and this cell type diversity is critical for increasing the computational capacity of neural circuits^7^. Cell type-specific differences in neuronal function are generally attributed to differences in electrical signal processing^6,7^, and diverse firing patterns emerge from differential expression and localization of ion channels^7,8^ as well as cell type-specific neuronal architectures^9,10^. However, chemical signal processing also shapes neuronal computation^11^. Calcium plays a particularly pivotal role in chemical signaling, regulating processes ranging from synaptic plasticity to synaptic vesicle fusion and neurotransmitter release, and neurons exhibit diverse calcium signals, from transient, local signals confined to single dendritic spines or axonal boutons to sustained, global signals that invade the entire neuron^12^. Cytoplasmic calcium fluctuations are generally attributed to influx across voltage gated channels in the plasma membrane (PM), but calcium signals do not directly reflect voltage signals^13^, indicating that additional mechanisms also regulate intracellular calcium signals. The endoplasmic reticulum (ER) is a major intracellular calcium store and, as such, is a likely candidate for shaping calcium signals that vary across neuronal cell types and subcellular compartments.

The ER is a dynamic network of cisternae and tubules that extend throughout dendrites and axons and form close contacts with the plasma membrane, mitochondria, and other organelles^14–16^. There is abundant *ex vivo* evidence that neuronal activation drives ER calcium responses^17–25^. In cultured neurons, the ER can act as a calcium sink^23,24^ or a calcium source^16,22^, taking up or releasing calcium upon stimulation. Moreover, the ER is not a passive calcium buffer. Like the plasma membrane, the ER is excitable: synaptic input triggers positive feedback loops in which calcium-and IP3-gated ER calcium channels amplify and propagate local calcium signals^12^. In principle, diverse ER calcium signals could emerge from differential expression and localization of these channels, as well as cell- and compartment-specific differences in ER architecture^26^. In addition, whereas excitation and propagation of electrical signals regulates calcium influx via voltage-gated ion channels in the plasma membrane, production and diffusion of reactive second messengers (e.g. IP3) regulates calcium efflux from the ER. ER signal processing therefore operates over long time scales (hundreds of milliseconds to minutes) and short distances (nanometers to microns) compared to signal processing at the plasma membrane, whereas voltage signals rapidly rise and fall (over millisecond time scales) and propagate over long distances (hundreds of microns to meters). Altogether, calcium signal processing by the ER has the capacity to expand the range of signal processing properties available to “the neuron,” thereby contributing to computational diversity across neuronal cell types. However, there are no direct measurements of stimulus-driven ER calcium signals in neurons *in vivo*, and the extent to which ER calcium signals vary across cell types and subcellular compartments in intact neural circuits *in vivo* is unknown.

In this work, we used *in vivo* two photon microscopy to image stimulus-evoked ER calcium signals in different neuronal cell types in the *Drosophila* visual system. The fly visual system is organized in a retinotopic, laminar fashion (Figure 1A)^27^. Photoreceptors in the retina synapse onto first-order interneurons in the first neuropil of the optic lobe (the lamina) which, in turn, synapse onto second-order interneurons in the second neuropil (the medulla). Second-order interneurons split into parallel processing streams that selectively respond to light increments (increases in luminance in the visual scene, or ON signals) or decrements (decreases in luminance, or OFF signals)^28–30^. ON and OFF second order interneurons synapse onto local motion detectors: T4 neurons in the ON pathway and T5 neurons in the OFF pathway^30^. Four classes of T4/T5 neurons selectively respond to motion in the four cardinal directions and project to one of four layers in the third neuropil (the lobula plate) based on their direction-selectivity; T4a/T5a selectively respond to local front-to-back motion and target layer one of the lobula plate, T4b/T5b respond to back-to-front motion and target layer two, and so on^31^. Lastly, T4/T5 neurons synapse onto wide-field lobula plate tangential cells (LPTCs), which detect specific patterns of global optic flow by integrating inputs from hundreds of T4/T5 neurons^32^. Extensively-studied LTPCs called HS (“Horizontal System”) neurons integrate input from T4a/T5a neurons in lobula plate layer one and thus selectively respond to global motion in the front-to-back direction^33–36^.

**Figure 1:**
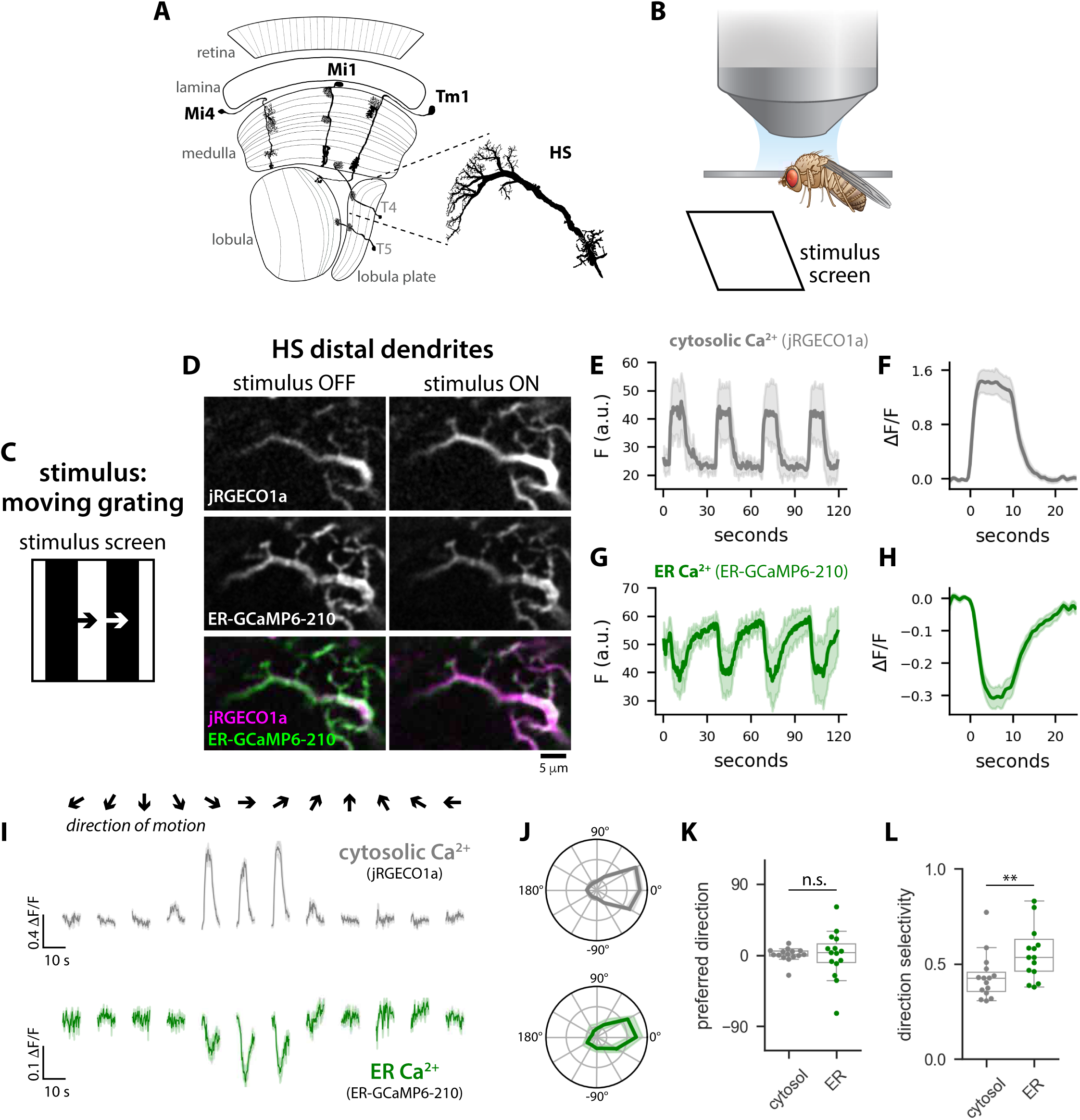
Global motion drives ER calcium release in HS dendrites. A: Schematic depiction of the *Drosophila* visual system. Example reconstructions of second order interneurons (Mi1, Mi4, and Tm1), local motion detectors (T4 and T4), and an HS neuron were adapted from Fischbach and Dittrich^35^. B: Experimental setup: *in vivo* two photon imaging of head-fixed flies with visual stimuli presented on a screen positioned in front of the fly. C: Visual stimulus: full contrast moving square wave gratings. D: Representative two photon images (average projections over a time series) of cytosolic calcium (jRGECO1a) and ER calcium (ER-GCaMP6-210) in HS distal dendrites before (stimulus OFF) and after (stimulus ON) the onset of motion. E,G: Raw cytosolic calcium (jRGECO1a, E) and ER calcium (G, ER-GCaMP6-210) responses to four bouts of global motion plotted over time; N=16 ROIs from one fly. F,H: Average change in cytosolic calcium (F) and ER calcium (H) averaged across all bouts of motion; N = 98 ROIs from 8 flies. I: Average cytosolic (gray) and ER (green) calcium responses to square waves gratings moving in different directions (arrows indicate direction of motion); N = 15 flies. J: Amplitude of cytosolic (gray) and ER (green) calcium responses to global motion in 12 different directions, normalized to the maximum response for each fly and plotted as a function of direction in polar coordinates. K-L: Box plots showing preferred direction (K) and direction selectivity index (L) for cytosolic and ER calcium responses; dots overlaid indicate average values for individual flies (N=15 flies).

Here, we leveraged *Drosophila* genetic tools to express an ER-targeted calcium reporter in four neuronal cell types in the visual system: HS neurons, two second-order ON neurons (Mi1 and Mi4), and one second-order OFF neuron (Tm1). By combining *in vivo* two photon imaging of ER calcium signals in head-fixed flies with visual stimulus presentation, we measured diverse stimulus-driven ER calcium responses that ranged from transient calcium uptake to sustained calcium release depending on the cell type, subcellular compartment, and stimulus.

## RESULTS

### Global motion drives ER calcium release in HS dendrites

There is abundant evidence that neuronal activation drives ER calcium release in dendrites *ex vivo*^16–18,20,22^. To determine if neuronal activation causes ER calcium release in dendrites *in vivo*, we used two photon microscopy to image cytosolic and ER calcium signals in *Drosophila* HS neurons while simultaneously driving neuronal activity with a visual stimulus (Figure 1, S1). There are three HS neurons per optic lobe, and each neuron has a large, elaborately branched dendrite that selectively responds to front-to-back global optic flow^33–36^. HS dendrites are easily accessible by *in vivo* two photon microscopy, and previous work has shown that preferred direction (PD) global motion (front-to-back) drives robust calcium signals in the cytosol of HS dendrites^35,36^. We used the GAL4/UAS system to express a low affinity, green fluorescent calcium reporter targeted to the ER lumen (ER-GCaMP6-210^23^) and a red fluorescent cytosolic calcium reporter (jRGECO1a^37^) in HS neurons. Then, we used two photon microscopy to image HS dendrites in head-fixed flies while simultaneously projecting moving square wave gratings on a screen positioned in front of the fly (Figure 1B-C). For each imaging z-plane, we presented four bouts of motion; for each bout, the square waving grating (λ = 30°) was stationary for 5 seconds, moved in the front-to-back direction (speed = 30°/s) for 10 seconds, and then remained stationary for another 15 seconds. To quantify cytosolic and ER calcium responses to these moving square wave gratings, we measured fluorescence intensities in regions of interest (ROIs) manually segmented from HS distal dendrites (Figure S1B). Consistent with previous work^35,36^, PD global motion drove large cytosolic calcium responses in HS distal dendrites, with cytosolic calcium levels rapidly increasing in response to the onset of motion and returning to baseline upon cessation of motion (Figure 1D-F, Figure S1C).

Strikingly, PD global motion also drove robust ER calcium responses, with calcium levels in the ER decreasing in response to motion before slowly returning to baseline (Figure 1D, G-H, Figure S1D). Stimulus-driven calcium response amplitudes varied across ROIs, even within the same imaging z-plane (Figure S1B-D), but there was no correlation between cytosolic and ER calcium response amplitudes (Figure S1E). However, we measured a slight correlation between ER calcium responses and dendrite branch thickness (Figure S1F), and ER calcium response amplitudes were significantly higher in thin, more distal dendrite branches (width < 1.4 µm) than in thicker, more proximal branches (width > 2 µm) (Figure 1G). This spatial variation in ER calcium response amplitudes may be due to subcellular patterning of ER calcium channels (i.e. RyR and IP3R), as observed in Purkinje cells^38^, and/or spatial variation in ER morphology, as recently measured in cultured rat hippocampal neurons^16^. Finally, we also found that the ER selectively responds to global PD motion. Global motion in non-preferred directions (e.g. vertical motion, or horizontal motion from back-to-front) failed to evoked calcium responses in either the cytosol or the ER (Figure 1I-L), consistent with previous measurements of HS direction-selectivity^35,36^. Local PD motion stimuli drove calcium responses in the cytosol but not in the ER (Figure S2). Thus, visual input drives ER calcium release in HS dendrites *in vivo* in a stimulus-selective fashion.

### Light increments drive compartment-specific ER calcium signals in Mi1 neurons

Our *in vivo* measurements of ER calcium release in HS dendrites (Figure 1) are consistent with *ex vivo* measurements of stimulus-driven ER calcium efflux in the dendrites of Purkinje cells^20^, hippocampal neurons^16^, and *Drosophila* mushroom body neurons^25^. In separate *ex vivo* experiments, neuronal activation has been shown to drive ER calcium uptake in the axons of hippocampal neurons^23,24^. Taken together, this body of work suggests that ER calcium handling is compartment-specific, with the ER acting as a calcium source in dendrites and a calcium sink in axons. However, reported differences in axonal versus dendritic ER responses could also be due to differences in experimental approach (e.g. neuronal activation via local glutamate uncaging^16^ versus field stimulation^23^), cell type-specific differences rather than compartment-specific differences, or some combination thereof.

To determine whether ER calcium handling in neurons is truly compartment-specific, we used our *in vivo* imaging setup to measure axonal and dendritic ER calcium responses to the same visual stimulus in the same cell type. HS axon terminals project to the lateral protocerebrum in the central brain, which is not optically accessible with our experimental setup. We therefore opted to image neurons upstream from HS, Mi1 neurons. Mi1 neurons are second-order interneurons that span the second neuropil of the optic lobe (the medulla) (Figure 2A) and form dense synaptic connections in four layers: M1, M5, M9, and M10^2,30^. Connections in M1 and M5 are mostly post-synaptic (∼75%) while M9 and M10 are mostly pre-synaptic (∼60% and ∼80% respectively)^39^. This mixed connectivity is common in *Drosophila* neurons^39^, and we classified subcellular compartments as either dendrites or axons based on the preponderance of connections (i.e. M1/M5 compartments are dendrites and M9/M10 compartments are axons). Mi1 neurons are ON neurons that depolarize in response to increases in luminance (light increments) and hyperpolarize in responses to decreases in luminance (light decrements)^13,28,29^. We therefore drove Mi1 activity by presenting alternating two second light and dark flashes that covered the entire visual stimulus screen (Figure 2B). We analyzed cytosolic and ER calcium responses in neurons with receptive field centers pointing at the stimulus screen (see Methods, Figure S3). In the cytosol, stimulus-locked calcium signals were similar in all subcellular compartments: calcium increased in response to light increments and decreased in response to light decrements in all compartments, although response amplitudes were significantly smaller in the dendritic compartments (M1/M5) than in the axonal compartments (M9/M10) (Figure 2C, S4C). In contrast, calcium responses in the ER were compartment-specific. In the dendrites, light increments caused calcium levels in the ER to decrease, consistent with ER calcium release (Figure 2D, S4D). In axons, the same stimulus had the opposite effect: light increments drove transient increases in ER calcium, consistent with ER calcium uptake (Figure 2D, S4D).

**Figure 2.**
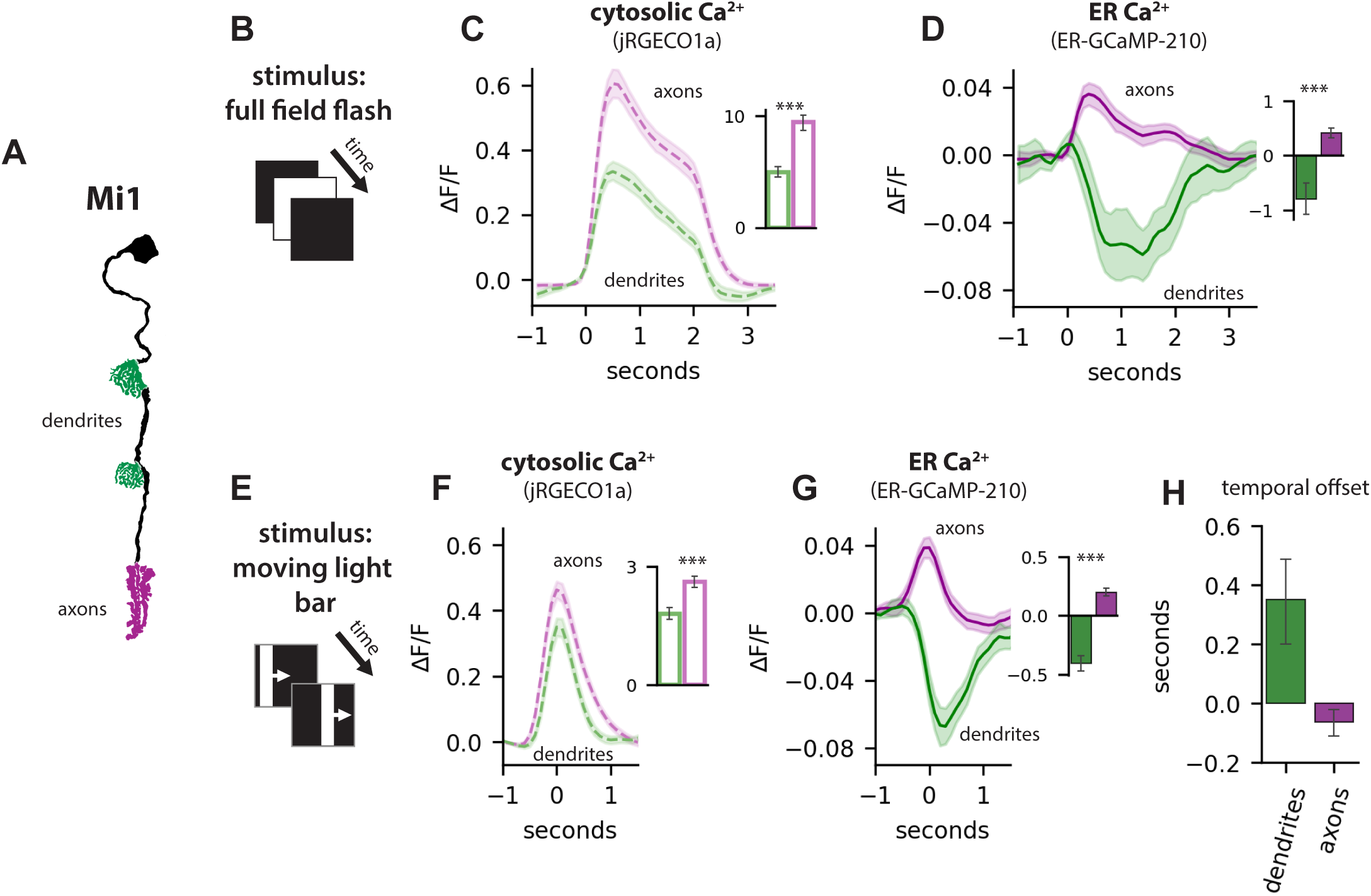
Compartment-specific ER calcium responses in Mi1 ON neurons. A: Mi1 neurons, with dendrites in green and axons in magenta. B: Full field flash stimulus. C-D: Average cytosolic calcium (C) and ER calcium (D) responses to full field flash stimuli in dendrites (green) and axons (magenta). Insets show average response amplitudes; N = 170 ROIs from 8 flies (dendrites) and 428 ROIs from 13 flies (axons). Asterisks indicate significant differences (Mann Whitney Test, p <0.05). E: Moving light bar stimulus. F-G: Average cytosolic calcium (F) and ER calcium (G) responses to moving light bars in dendrites (green) and axons (magenta). Insets show average response amplitudes; N = 154 ROIs from 8 flies (dendrites) and 391 ROIs from 19 flies (axons). H: Time to peak ER calcium responses, calculated relative to the peak cytosolic response.

To investigate whether compartment-specific ER calcium signals in Mi1 neurons are conserved in response to a different visual stimulus, we imaged ER and cytosolic calcium in axons and dendrites while presenting a light bar moving across a dark background (Figure 2E). Mi1 receptive fields have a weak inhibitory surround as well as an excitatory center^40^. Whereas full field light increments simultaneously activate the center and the surround, moving light bars sequentially activate the surround and then the center as the bar moves across the visual scene. Moving bars also stimulate neighboring neurons in succession (Figure S3), so we mutually aligned ER calcium responses from a large population of cells based on cytosolic calcium response peaks (Figure 2F). Consistent with responses to widefield light increments, we measured a delayed decrease in ER calcium in dendrites and a rapid, transient increase in ER calcium in axons (Figure 2F-H). Altogether, these results demonstrate that the ER can simultaneously act as a calcium source and a calcium sink in different subcellular compartments of the same neuron, releasing calcium in dendrites while taking up calcium in axons.

### Visual stimulus-driven ER calcium responses are cell type-specific

Next, we wished to determine if compartment-specificity — i.e. stimulus-driven release in the dendrites and uptake in the axons — is conserved across cell types. To that end, we measured visual stimulus-driven cytosolic and ER calcium responses in two additional second-order interneurons: Mi4 and Tm1 neurons. Like Mi1, Mi4 and Tm1 respond to changes in luminance in the visual scene, but the three cell types have distinct morphologies^2^, contrast preferences^28,29,41^, spatiotemporal filtering properties^40^, transcriptional profiles^42–44^, and synaptic inputs^39^ (Figure S5A-C). Thus, Mi4 and Tm1 are similar enough to Mi1 to respond to the same simple visual stimuli (e.g. alternating widefield light increments and decrements), but different enough to test whether compartment-specific ER calcium responses are conserved across diverse neuronal cell types. In addition, three major molecular players mediate ER calcium handling: two ER calcium channels (ryanodine receptors (RyR) and IP3 receptors (IP3R)) allow calcium release from the ER and the sarcoplasmic/ER calcium ATPase (SERCA) pumps calcium into the ER in an ATP-dependent fashion^12^. In Mi1, the ER exhibits transient calcium fluctuations in response to visual input, with calcium decreasing in dendrites and increasing in axons before returning to baseline (Figure 2D). This suggests that synaptic input transiently increases RyR- and/or IP3R-dependent calcium efflux relative to SERCA uptake in dendrites and vice versa in axons. Interestingly, Mi1, Mi4, and Tm1 express significantly different levels of RyR and IP3R, with higher expression of IP3R in Mi4 and higher expression of RyR in Tm1 (Figure S5D). We were therefore curious to see whether Mi4 and Tm1 exhibit compartment-specific ER calcium release in dendrites and uptake in axons, like Mi1, or if higher expression of IP3R (in Mi4) or RyR (in Tm1) tips the calcium influx/efflux balance towards efflux in all compartments.

Like Mi1, Mi4 has distinct subcellular compartments (Figure 3A, S5B): dendrites that arborize in medulla layers M2-M5 and are primarily (∼70%) post-synaptic and axons in M8 and M10 that are primarily presynaptic (60% and 70% respectively)^2,39^. Mi4 is also an ON neuron that depolarizes in response to light increments^41^, but Mi1 and Mi4 have different response kinetics: Mi1 responds to increments in a transient fashion, whereas Mi4 responses are more sustained^40^. To determine whether Mi4 ER calcium responses exhibit subcellular compartmentalization, we measured cytosolic and ER calcium responses to widefield light increments (Figure 3C-D) and moving light bars (Figure 3E-G). In the cytosol, both stimuli drove large, sustained increases in cytosolic calcium levels in all subcellular compartments (Figure 3C,F). In Mi4 dendrites, widefield light increments drove larger decreases in ER calcium levels than in Mi1 dendrites, consistent with higher expression of IP3R in Mi4 neurons (Figure 3D). In the axons, widefield light increments drove slight ER calcium uptake in M10, similar to Mi1, and no significant response in M8, indicating that Mi4 ER calcium responses to wide field light increments are compartmentalized in a similar fashion to Mi1, with the dendritic ER acting as a net calcium source and the axonal ER (in M10) acting as a slight calcium sink (Figure 3D).

**Figure 3.**
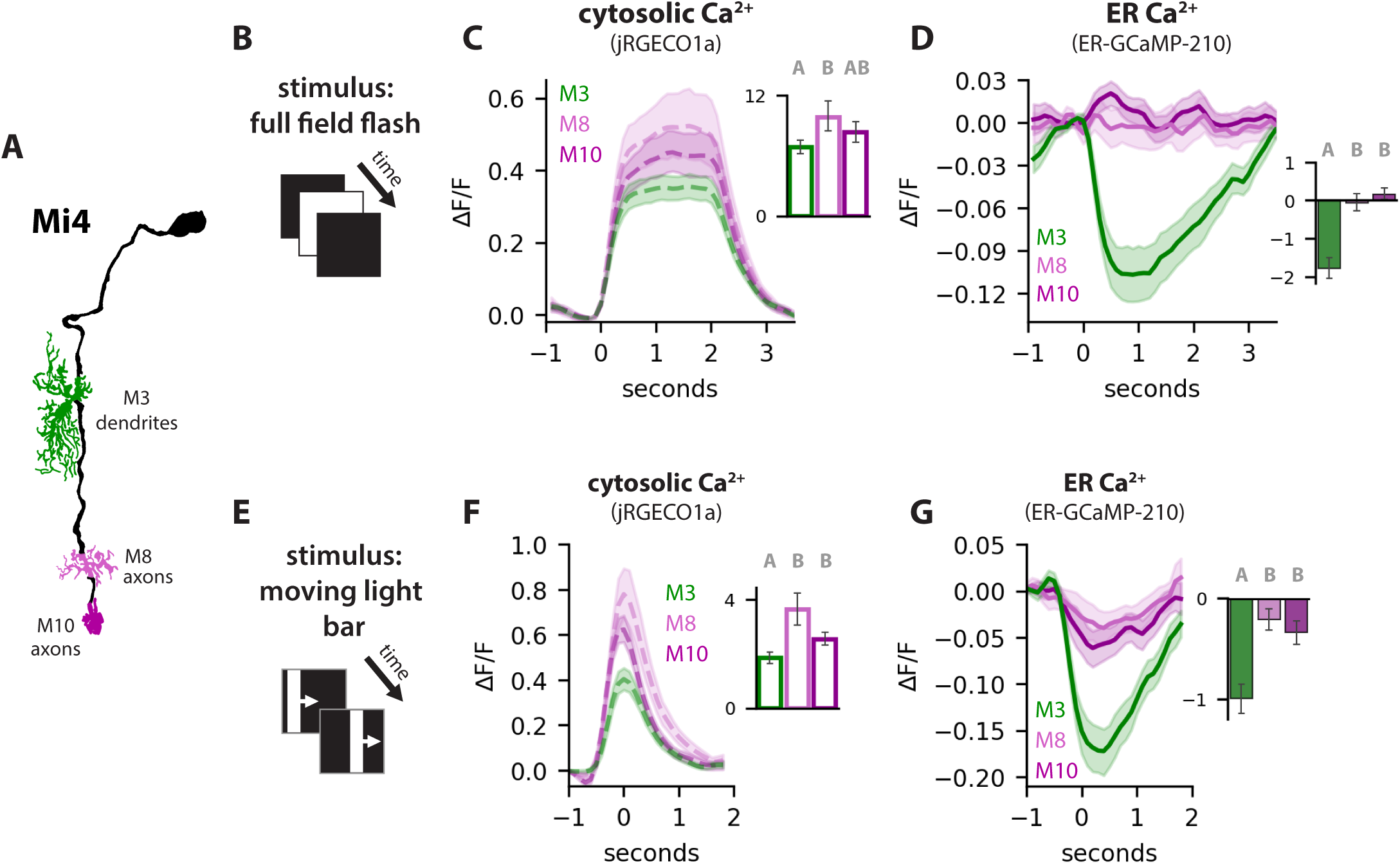
Motion stimuli drive calcium efflux from all subcellular compartments in Mi4 ON neurons. A: Mi4 neuron with dendrites in M3 in green and axons in M8 and M10 in shades of magenta. B: Full field flash stimulus. C-D: Average cytosolic calcium (C) and ER calcium (D) responses to full field flash stimuli. E: Moving light bar stimulus. F-G: Average cytosolic calcium (F) and ER calcium (G) responses to moving light bars. Insets show average response amplitudes; N = 228 ROIs from 7 flies (M3 dendrites), 127 ROIs from 9 flies (M0 axons), and 154 ROIs from 9 flies (M10 axons). Letters on inset bar plots indicate groups that are significantly different (Kruskal-Wallis with post-hoc Dunn’s test).

Strikingly, we also found that the sign of the ER response in Mi4 axons (i.e. net uptake versus net release) varied as a function of stimulus type. Like Mi1, the Mi4 receptive field has an inhibitory surround and an excitatory center^40^. Whereas widefield light increments, which simultaneously activate the center and the surround, drove either no ER response (in M8) or slight ER calcium uptake (in M10) (Figure 3D), moving light bars, which sequentially activate the surround followed by the center, drove significant ER calcium release in all subcellular compartments, including axons in M8 and M10 (Figure 3G). Thus, unlike in Mi1, the axonal ER in Mi4 can switch from a slight calcium sink to a calcium source depending on the visual stimulus. In addition, motion-selective ER calcium release in Mi4 axons is consistent with the idea that the ER implements sequence-dependent coincidence detection, as previously proposed based on *ex vivo* measurements in Purkinje cells^45^ and *Drosophila* mushroom neurons^25^.

Having measured compartment- and stimulus-specific ER calcium responses in Mi4, we next measured ER responses in Tm1 neurons (Figure 4). Whereas Mi4 expresses high levels of IP3R, which is gated by coincident binding of IP3 and calcium, Tm1 expresses high levels of RyR, which is gated by calcium alone. We therefore expected that stimulus-driven increases in cytosolic calcium would be associated with rapid calcium release from the ER in Tm1 dendrites, and perhaps in Tm1 axons as well. Tm1 has three distinct compartments (Figure 4A, S5B): dendrites (∼95% postsynaptic) in medulla layer M1, axons (∼80% pre-synaptic) in the first layer of the third neuropil of the visual system, the lobula, and a “mixed” connectivity compartment in medulla layer M9 that contains equal numbers of pre- and post-synaptic connections (∼50% presynaptic and 50% postsynaptic)^2,42^. In contrast to Mi1 and Mi4, Tm1 is an OFF neuron that depolarizes in response to light decrements and hyperpolarizes in response to increments^28,29^. Consistent with this, we found that light decrements drove rapid increases in cytosolic calcium in all subcellular compartments of Tm1 (Figure 4B). However, ER calcium responses in Tm1 were entirely different from ER responses in Mi1 or Mi4. Despite driving increases in cytosolic calcium, light decrements drove only a small, transient reduction in ER calcium in the mixed connectivity compartment of Tm1 (in M9) and had no significant effect on ER calcium levels in either the dendrites (in M2) or the axons (in Lob1) (Figure 4C). Even more surprising, despite causing cytosolic calcium to decrease in all compartments, light increments drove significant, transient ER calcium uptake in in M9 and slight (but not significant) uptake in dendrites (M2) and axons (Lob1). It is unclear why the relatively high expression of RyR in Tm1 does not result in robust visual stimulus-driven ER calcium release in any subcellular compartment. It may be that stimulus-driven increases in ER calcium efflux are balanced by increased ER calcium uptake via SERCA, resulting in little to no net change in ER calcium levels. Regardless of the molecular mechanisms underlying ER calcium signals (or lack thereof) in Tm1 neurons, these results demonstrate that visual stimulus-driven ER calcium signals vary across cell types.

**Figure 4.**
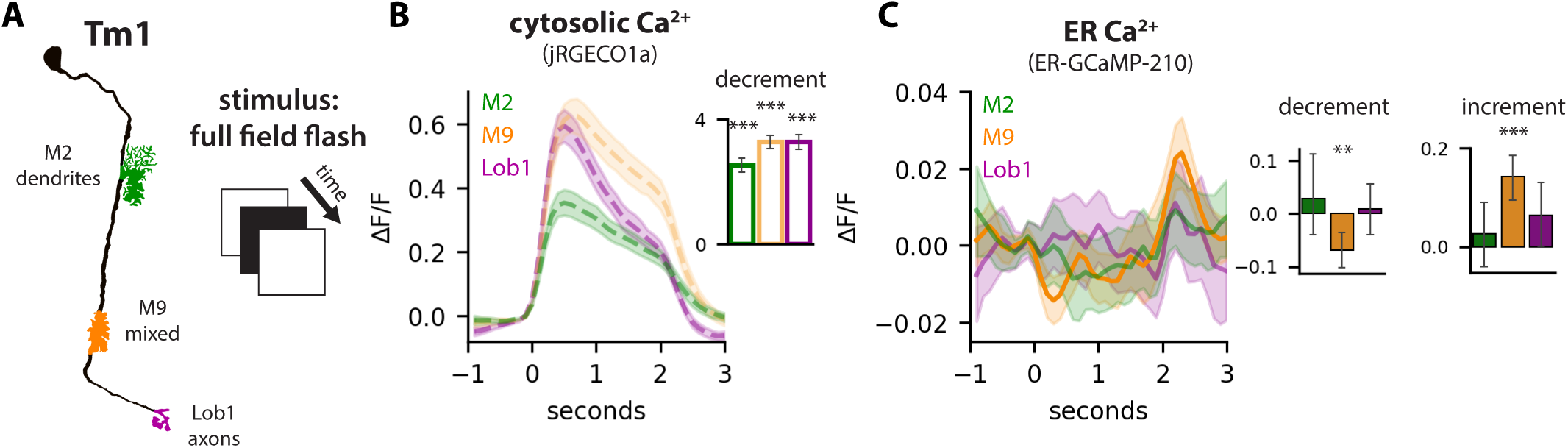
Light increments drive transient ER calcium uptake in Tm1 OFF neurons. A: Tm1 neuron with dendrites in M2 in green, mixed connectivity compartment in M9 in yellow, and axons in the lobula in magenta. B: Average cytosolic calcium responses to full field flash stimuli; bar plot shows averages response amplitudes for light decrements. C: Average ER calcium responses to full field flash stimuli; bar plots show averages response amplitudes for light decrements (left) and light increments (right). Asterisks indicate average values that are significantly different from zero (sign test, p < 0.05); N = 123 ROIs from 9 flies (M2), 189 ROIs from 10 flies (M9), and 167 ROIs from 9 flies (Lob1).

### IP3R is necessary and sufficient for visual stimulus-driven ER calcium release

Having measured ER calcium signals that vary as a function of cell type, subcellular compartment, and visual stimulus, we set out to explore the molecular mechanism underlying visual stimulus-driven ER calcium responses. First, we turned our attention back to HS dendrites, which exhibit ER calcium release in response to global motion. There is no correlation between cytosolic and ER calcium response amplitudes in HS dendrites (Figure S1E), suggesting that an additional signal, other than cytosolic calcium (e.g. IP3), regulates ER calcium release. We therefore hypothesized that IP3R mediates ER calcium release in HS dendrite. To test this hypothesis, we knocked down (via RNAi) or overexpressed IP3R in an HS-specific fashion and measured the effects of ER calcium responses to PD global motion (Figure 5A). IP3R knockdown abolished ER calcium responses, and IP3R overexpression significantly increased response amplitudes (Figure 5B-C). Thus, in HS dendrites, IP3R gates ER calcium release in response to global motion.

**Figure 5.**
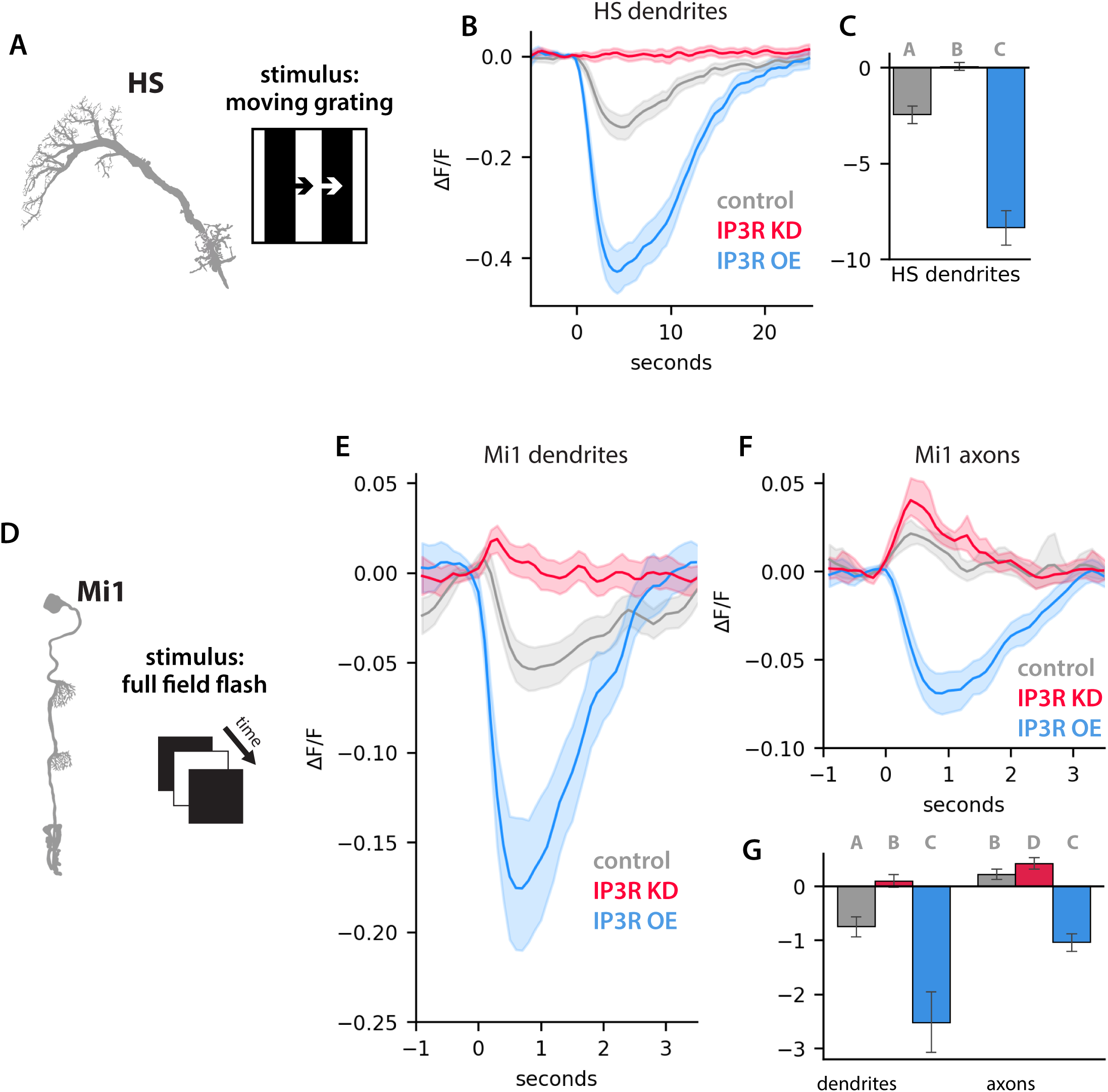
IP3R is necessary and sufficient for visual stimulus-driven ER calcium release in Mi1 neurons. A: Schematics indicating cell type/compartment (HS dendrites) and visual stimulus (moving square wave gratings). B-C: ER calcium (ER-GCaMP6-210) signals in HS dendrites plotted over time (B) and average responses amplitudes (C). N = 113 ROIs from 4 flies (control, gray), 112 ROIs from 4 flies (IP3R knockdown, red), and 149 ROIs from 5 flies (IP3R overexpression, blue); letters indicate groups that are significantly different (Kruskal Wallis with post-hoc Dunn’s test. D: Schematics indicating cell type (Mi neurons) and visual stimulus (full field flashes). E-G: ER calcium (ER-GCaMP6-210) signals in Mi1 dendrites and axons plotted over time (E-F) and average responses amplitudes (G). N = 227 ROIs (10 flies) in control dendrites, 205 ROIs (6 flies) in IP3R KD dendrites, 62 ROIs (3 flies) in IP3R dendrites, 504 ROIs (17 flies) in control axons, 407 ROIs (13 flies) in IP3R KD axons, and 230 ROIs (9 flies) in IP3R axons. Letters over the bar plots indicate significantly different groups (Kruskal Wallis test with post-doc Dunn’s test, p < 0.05).

Finally, we explored the role of IP3R in subcellular compartmentalization of ER calcium responses in Mi1 neurons. We reasoned that simultaneous ER calcium release in dendrites and uptake in axons could be due to enrichment of IP3R (and/or stimulus-driven production of IP3) in Mi1 dendrites compared to axons, and we knocked down or overexpressed IP3R in an Mi1-specific fashion and measured the effects on ER calcium responses to widefield light increments (Figure 5D). Strikingly, both manipulations (IP3R knock down and overexpression) disrupted compartment-specificity (Figure 5E-G). IP3R knockdown switched the dendritic ER from a calcium source to a calcium sink (Figure 5E), such that visual input drove ER calcium uptake in all compartments. IP3R overexpression switched the axonal ER from a calcium sink to a calcium source (Figure 5F), such that visual input drove calcium release from the ER in all compartments. In addition, compared to controls, IP3R knockdown increased the amplitude ER calcium uptake in axons, and IP3R overexpression increased the amplitude of ER calcium release in dendrites (Figure 5E-G). Altogether, these results are consistent with a model in which visual input increases calcium flux across the ER membrane, with net efflux in dendrites due to enrichment of IP3R relative to SERCA and vice versa in axons.

## DISCUSSION

Our results demonstrate that visual input drives diverse ER calcium signals in *Drosophila* neurons *in vivo.* We found that the timing, amplitude, and sign (uptake versus release) of these ER calcium signals vary as a function of cell type (e.g. substantial ER calcium release in Mi4 dendrites versus no significant response in Tm1 dendrites), subcellular compartment (e.g. slow calcium release in Mi1 dendrites versus fast, transient uptake in Mi1 axons), and spatial extent of the visual stimulus (e.g. sustained calcium release in response to global but not local motion stimuli in HS dendrites). These diverse ER calcium signals are not simply a reflection of context-specific differences in cytosolic calcium signals. Visual stimulus-driven increases in cytosolic calcium were variously associated with increases, decreases, or no net change in ER calcium, and in one cell type (Tm1), decreases in cytosolic calcium were associated with transient ER calcium uptake. Thus, rather than acting as a passive calcium buffer, the ER actively processes calcium signals in a cell type and compartment-specific fashion.

Cell type diversity in electrical signal processing relies on differential expression and subcellular patterning of ion channels in the plasma membrane^7,8^. Similarly, cell- and compartment-specific ER calcium signals likely depend on differential expression and subcellular patterning of ER calcium channels. In Mi1 neurons, we found that proper expression of IP3R is necessary for compartment-specific ER calcium responses to visual input: IP3R knockdown abolished stimulus-driven calcium efflux from the dendritic ER, and IP3R overexpression was sufficient to switch the axonal ER from a calcium sink to a source. These IP3R-dependent, compartment-specific ER responses may be the result of subcellular patterning of IP3R, with IP3R enriched in dendrites relative to axons. Alternatively, if IP3R is densely distributed throughout dendrites and axons, as previously observed in Purkinje cells via immunogold labelling of IP3R^46^, dendrite-specific ER calcium release could be achieved by dendrite enrichment of stimulus-driven IP3 production. Various Gαq-coupled GPCRs in *Drosophila*^47^, including metabotropic neurotransmitter receptors as well as neuromodulator and neuropeptide receptors, could induce IP3 production in response to synaptic (or extra-synaptic) input onto Mi1 dendrites. It is difficult to measure localization patterns in specific neurons within a densely-packed neuropil. However, a genetic approach for cell type-specific endogenous tagging recently revealed subcellular patterning of neurotransmitter receptors in *Drosophila* neurons, including cell type-specific differences in the degree of enrichment in dendrites versus axons^48^; a similar approach may reveal cell type-specific subcellular patterning of ER calcium channels.

In addition to mediating different ER responses between dendrites versus axons, the spatial arrangement of ER calcium channels and/or neurotransmitter receptors within dendrites or axons may confer stimulus specificity, with the ER selectively amplifying calcium signals in response to specific synaptic input patterns. In HS dendrites, we found that global motion, which activates the entire dendrite, drives IP3R-dependent ER calcium release, but local motion, which activates short segments of individual dendritic branches, does not. In contrast, local glutamate uncaging has recently been shown to induce RyR-dependent ER calcium release in cultured rat hippocampal neurons^16^. ER excitability (or lack thereof) by local synaptic input may depend on the proximity of ER calcium channels to calcium channels in the plasma membrane. In hippocampal neurons, RyR channels cluster with voltage-gated calcium channels at ER-PM contact sites, which likely exposes RyR to high local concentrations of calcium upon local spine activation^16^. In HS dendrites, IP3R may be situated at a remove from the plasma membrane, requiring broader dendrite activation to generate sufficiently high concentrations of IP3 and calcium to gate ER calcium release. Alternatively, selective excitation of IP3R-dependent calcium by global versus local (or strong versus weak) synaptic input may depend on the spatial arrangement of metabotropic and ionotropic neurotransmitters at the synapse. In hippocampal neurons, ionotropic glutamate receptors (AMPAR) are positioned in the middle of the postsynaptic density, immediately adjacent to synaptic release sites in the presynaptic terminal, but metabotropic glutamate receptors (mGluR) localize to the periphery of the presynaptic density^49^. As a result, weak synaptic input (ie low levels of glutamate release from the presynaptic neuron) engage AMPA receptors, but larger, more sustained synaptic input (more glutamate) is required to engage mGluR at the periphery of the postsynaptic density. Similarly, transient local motion stimuli may sufficient to engage calcium-permeable ionotropic neurotransmitter receptors in HS dendrites (e.g. nAChR) while sustained global motion may be necessary to engage Gαq-coupled metabotropic neurotransmitter receptors (e.g. mAChR-A) and excite IP3R-dependent ER calcium release.

In addition to setting a threshold for ER excitation based on synaptic input strength, fine spatial patterning of ER calcium channels and/or neurotransmitter receptors could also mediate ER selectivity based on synaptic input timing. In Mi4 axons, we found that ER calcium release is motion-selective: sequential activation of Mi4 inputs by a moving light bar triggered ER calcium release, whereas simultaneous activation induced slight ER calcium uptake. Mi4 expresses high levels of IP3R, and whereas IP3 binding to IP3R exposes a calcium binding site necessary for channel opening^50^, calcium regulates IP3R activity in a biphasic fashion: low calcium concentrations activating and high concentrations inhibiting channel activity^51^. Mi4 axons receive direct synaptic input, and motion-selective ER calcium release may be the result of sequential engagement of spatially segregated metabotropic and ionotropic neurotransmitter receptors within the axonal arbor, resulting in IP3 production followed by cytosolic calcium transients and thus more effective activation of IP3R. Similarly, ER calcium release via IP3R has been proposed to mediate sequence-dependent coincidence detection in Purkinje cells^52^ and *Drosophila* mushroom body neurons^25^.

Finally, a large body of *ex vivo* evidence implicates ER calcium handling in various aspects of neuronal function, including plasma membrane excitability^53^, synaptic plasticity^20,21,25,45,52,54–56^, and synaptic transmission^23,24,57,58^. In addition, genetic augmentation of ER calcium release into the cytosol has been shown to enhance dendritic feature selectivity and place field stability in hippocampal pyramidal neurons^59^, indicating that ER calcium handling can tune neuronal input-output functions *in vivo*. Our results demonstrate that visual input elicits diverse ER calcium signals that vary as a function of cell type and subcellular compartment in *Drosophila* neurons. Taken together, this body of work suggests that cell type diversity in ER-based calcium signal processing has the potential to expand the computational capacity of neural circuits.

## ACKNOWLEDGEMENTS

This work was supported by the NIH (R01EY036445 to ELB and F31EY035165 to KCD) and the NSF (grant number 2227609 to ELB).

## AUTHOR CONTRIBUTIONS

Conceptualization, ELB and KCD; Investigation, KCD, KW, LL, and ELB; Software KRK and ELB; Formal Analysis, KCD, KRK, and ELB; Writing — Original Draft, KCD; Writing — Review and Editing; KCD and ELB; Supervision, ELB.

## METHODS

### *Drosophila* strains and husbandry

All flies were maintained in vials on a standard molasses food at 25°C and 60% humidity in a 12h light/dark cycle. Crosses were flipped onto fresh food every 3 days. Progeny were imaged 5-7 days after eclosion. Drosophila strains used in this study include:

| Strain | Bloomington ID |
| --- | --- |
| Wild-type Canton S | BDSC 64349 |
| Wild-type isogenized | BDSC 99750 |
| R27B03-GAL4 | HS driver, BDSC 49211 |
| R19F01-GAL4 | Mi1 driver, BDSC 48852 |
| R48A07-AD/ R13F11-DBD | Mi4 split GAL4, BDSC 86853 |
| R74G01-GAL4 | Tm1 driver, BDSC 39868 |
| UAS-IP3R RNAi | BDSC 51686 |
| UAS-IP3R OE | BDSC 30742 |
| UAS-jRGECO1a | BDSC 63795 |
| UAS-ER-GCaMP-210 | BDSC 83294 |

### In vivo imaging

Female flies were cold anaesthetized and positioned in a keyhole shaped hole in a thin metal shim. Flies were positioned so that the eye sat below the shim and the back of the head was flat and above the shim. Each fly was glued in place using a UV-cured glue (Bondic). Using fine forceps, a small hole was dissected in the cuticle of the fly and fat and trachea were removed to expose the brain. The brain was submerged in a sugar saline solution (103mM NaCl, 3mM KCl, 5mM TES, 1mM Na_2_PO_4_, 26mM NaHCO_2_, 4mM MgCl_2_, 1.5mM CaCl_2_, 10mM trehalose, 10mM glucose and 7mM sucrose) that was bubbled with carbogen to oxygenate and regulate pH. Mi1, Mi4, and Tm1 neurons were imaged using an integrated confocal and 2-photon microscope (Leica SP8 CSU MP Dual) with a 25x 1.0 NA water immersion objective (Leica). For 2-photon imaging, a tunable MP laser (Insight X3, Spectra-Physics) tuned to 920nm was used to excite ER-GCaMP6-210 and a fixed 1045nm laser was used to excite jRGECO1a. Emitted photons were collected using HyD detectors (Leica). 200x200 images were collected at a frame rate of 10Hz, 15x digital zoom and bidirectional scanning. HS dendrites were imaged using an Olympus FV1000 microscope equipped with a 25x 1.05 NA water immersion objective (Olympus). A Spectra-Physics Mai Tai Ti:Sapphire laser tuned to 1000nm was used simultaneously excite ER-GCaMP6-210 and jRGECO1a. Emitted photons were collected using a 520/40nm emission filter. 256x256 images were collected at a frame rate of 5Hz, with 12x digital zoom, and bidirectional scanning. Imaging time per fly never exceeded 2 hours.

### Visual stimulus presentation

Visual stimuli were generated using PsychoPy and presented on a flat white screen (Da-Lite Dual-Vision Vinyl) using a digital light projector (DLP LightCrafter, Texas Instruments). The stimulus screen spanned ∼60° of the fly’s visual field in both the horizontal and vertical directions. The stimulus was filtered using a 472/30 nm bandpass filter (Semrock) before projection to avoid detection of stimulus light by the microscope. Voltage signals from the imaging software were relayed to PsychoPy using a LabJack device, synchronizing the stimulus and imaging frames. Global motion stimuli used to drive HS responses were full-contrast square wave gratings (λ = 30°) that spanned the entire stimulus screen and moved at 30°/s in the front-to-back direction with respect to the fly’s eye for 10 seconds; bouts of motion were interspersed by 20 second presentations of static gratings. To measure HS direction selectivity, square wave gratings moving in one of 12 different directions were presented in a randomized order; 2 second bouts of motion were interspersed with 4 seconds of static grating presentation. Local motion stimuli used to drive local calcium responses in HS dendrites were 13.5° tall moving light edges that appeared on a black background and expanded 13.5° in the front-to-back direction over 0.5 seconds (speed = 27°/s) before disappearing. Full field flash stimuli used to drive responses in second-order interneurons were alternating full contrast light increments and decrements that spanned the entire stimulus screen; two seconds elapsed between each contrast step. Moving light bar stimulus, used to drive Mi1 and Mi4 responses, was a white bar (width = ∼15°) that moved across a black screen in four directions (up, down, right, and left) at ∼15°/s.

### Image analysis

Visual stimulus-driven calcium responses were analyzed using custom-written Python code. Pystackreg was used to align time series images in X and Y. Aligned images were binned by stimulus type (i.e. dark vs. light screen) and then by time (start of each stimulus epoch), and fluorescent intensities in regions of interest (ROIs) were extracted from binned images based on binary masks manually generated in Fiji. The average fluorescent intensity was calculated over time for each ROI and ΔF/F was calculated as (F_t_-F_0_)/F_0_ where F_0_ was the average baseline fluorescent intensity, calculated from the average fluorescence intensity within each ROI in the 1 second (for HS) or 250ms (for second-order interneurons) prior to stimulus onset (the onset of motion for HS and the light increment/decrement for ON/OFF second-order interneurons). To calculate response amplitudes for square wave grating, moving dark edge, or full flash flashes, ΔF/F values for each ROI were summed over the entire period of motion (for moving gratings or expanding dark edges) or the entire 2 second light or dark period (for full field flashes). Direction selectivity index (Figure 1L) was calculated by dividing the amplitude of the response to the preferred direction of motion by the sum of the responses to all 12 directions of motion. Moving bar responses for all ROIs corresponding to the same cell type/compartment were mutually aligned in time by cross-correlation of each individual jRGECO1a response with the average jRGECO1a response; responses were aligned in an iterative fashion, such that the average response was recalculated after each round of alignment, over 20 rounds of alignment. For each ROI, the temporal offset calculated based on the jRGECO1a response was applied to the ER-GCaMP6-210 response. To calculate moving bar response amplitudes, ΔF/F responses were summed over a 0.6 second time window centered around the peak response in the average trace for each cell type/compartment. For second-order interneurons, cells with receptive field centers off the visual stimulus screen (Figure S3) were excluded from subsequent analysis. For full field flashes, “off screen” neurons were removed by excluding ROIs with jRGECO1a response amplitudes below an arbitrary threshold (based on responses to light increments for Mi1/Mi4 and light decrements for Tm1). For moving light bars, off screen neurons were removed by excluding ROIs that failed to exhibit jRGECO1a responses to bars in all four directions of motion.

### Statistical Analysis

Datasets were tested for normality using the Shapiro-Wilk test. For non-normal data, single comparisons were performed using the Mann Whitney test and multiple comparisons were performed using the Kruskal-Wallis test followed by the Dunn’s test with a Bonferroni correction. All statistical tests were performed using the stats module in SciPy. The statistical test used for each dataset is indicated in the corresponding figure legend.

