## Supplemental Figures for "Visual input drives diverse ER calcium signals in neurons *in vivo*"

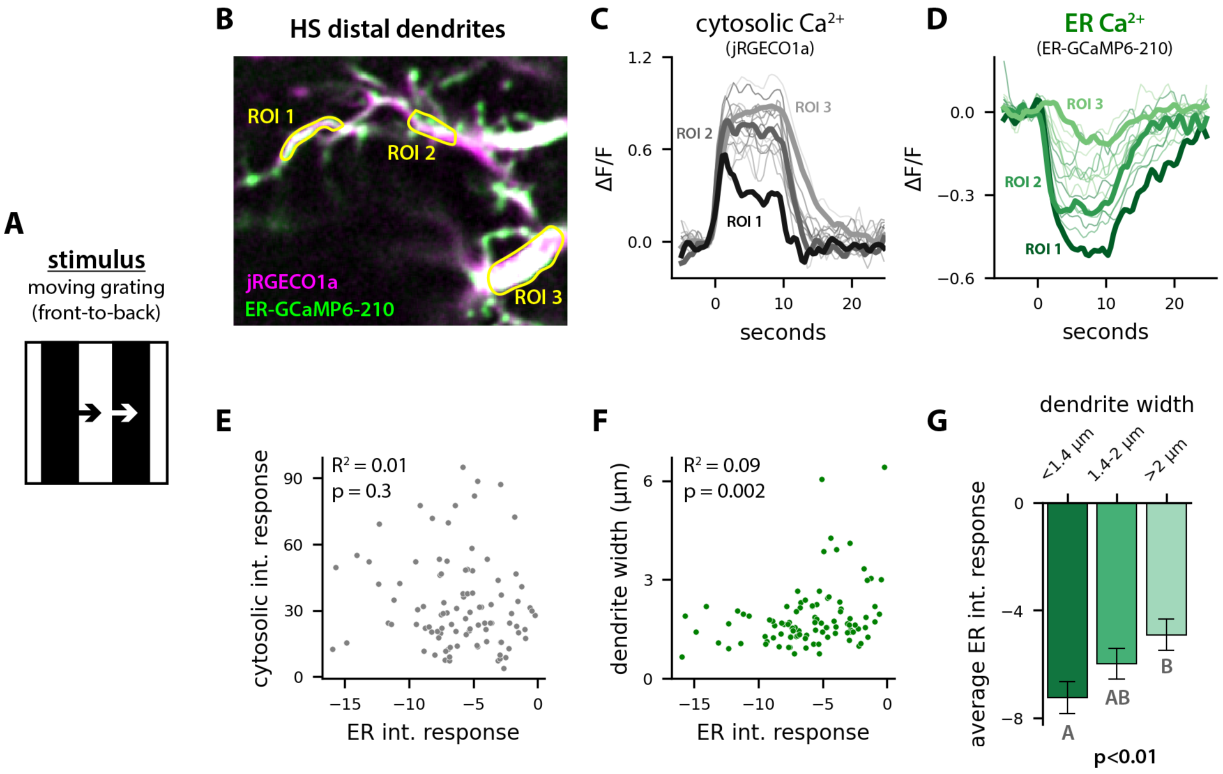


**Figure S1: Spatial variation in ER calcium response amplitudes in HS dendrites.** A: Visual stimulus: full contrast moving square wave gratings. B: Average projection (over a time series) of cytosolic (jRGECO1a, magenta) and ER (ER-GCaMP6-210, green) calcium signals in HS distal dendrites. Example regions of interest (ROIs) are indicated in yellow. C-D: Average change in cytosolic calcium (C) and ER calcium (D) in 15 ROIs extracted from dendrites shown in B. E-F: ER calcium response amplitudes plotted versus cytosolic calcium response amplitudes (E) or dendrite width (F); N = 98 ROIs from 8 flies. G: Bar plots indicating average ER calcium response amplitudes in ROIs binned by dendrite width; error bars are standard error of the mean and letters indicate groups that are significantly different from each other (Kruskal-Wallis with post-hoc Dunn's test, $p < 0.05$).

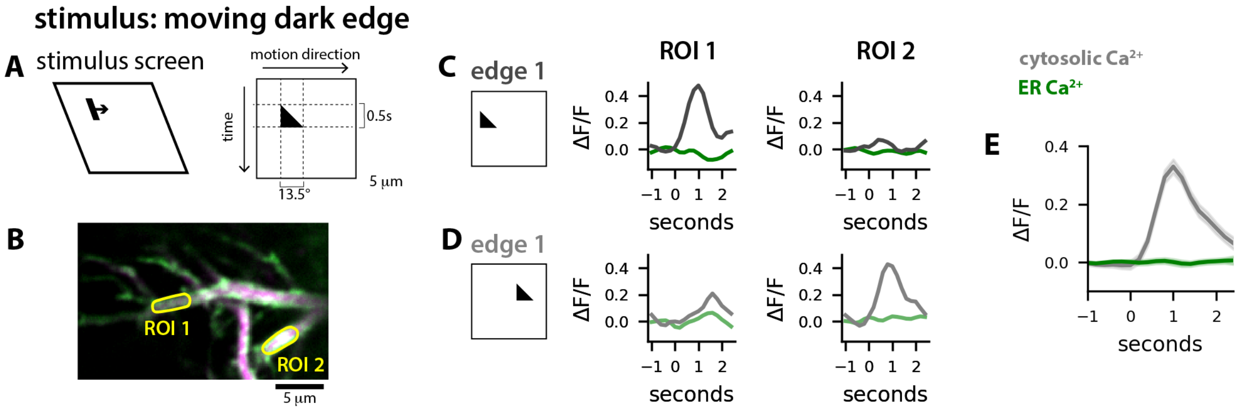


**Figure S2: Local motion does not drive ER calcium release in HS dendrites.** A: Visual stimulus screen (left) and a space-time plot (right) depicting the local motion visual stimulus. A single moving dark edge expanded in the preferred direction (PD) over 13.5° in 0.5 seconds (speed = 27°/s) before disappearing. B: Average projection (over a time series) of cytosolic (jRGECO1a, magenta) and ER (ER-GCaMP6-210, green) calcium signals in HS distal dendrites. Example regions of interest (ROIs) are indicated in yellow. C-D: Cytosolic (gray) and ER (green) calcium responses in the two ROIs indicated in B to local motion stimuli presented at two different points in space on the visual stimulus screen. Note that edge 1 drove local cytosolic calcium responses in ROI 1 (C) whereas edge 2 drove local cytosolic calcium responses in ROI 2 (D). E: Average cytosolic calcium (gray) and ER calcium responses to local moving dark edges; N = 58 ROIs from 6 flies.

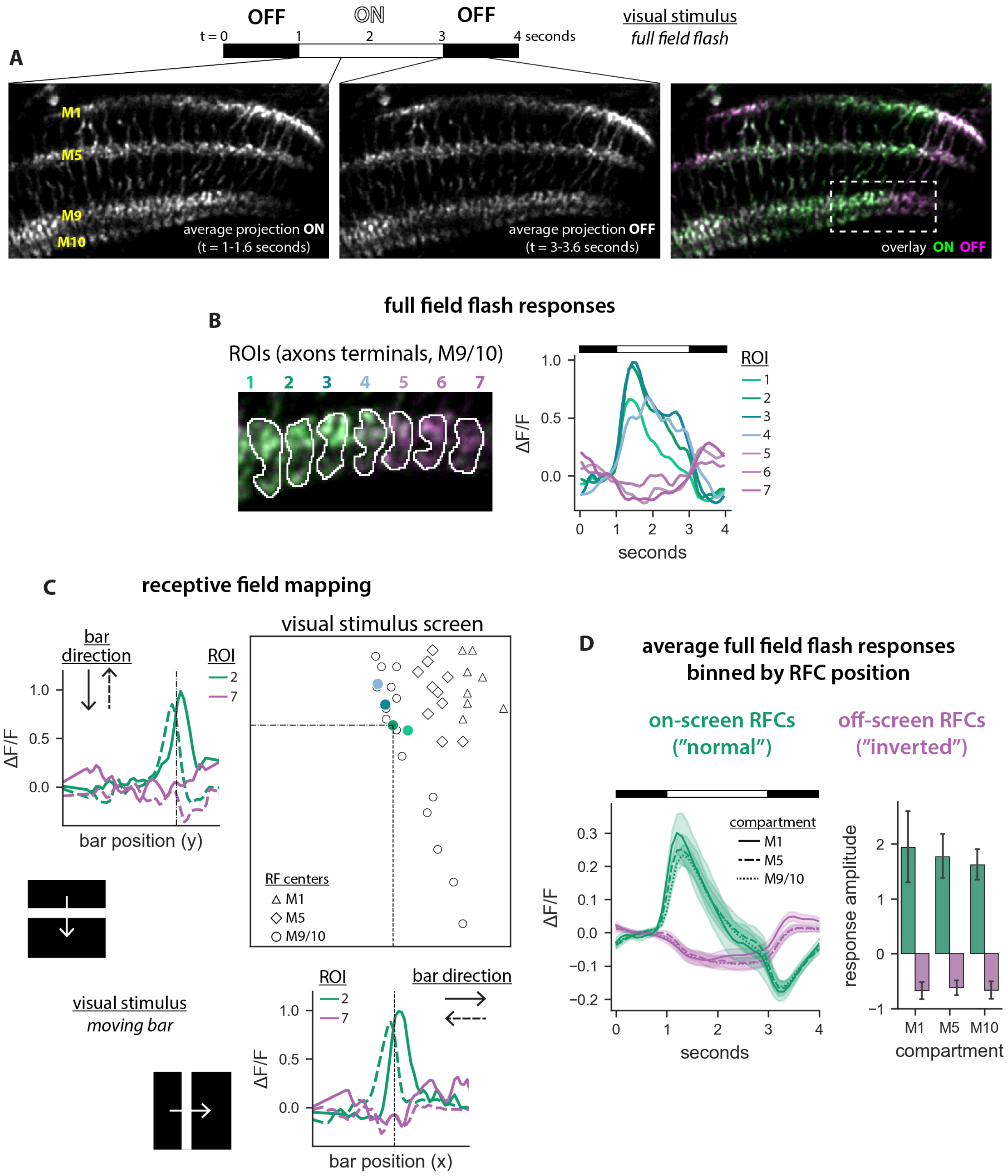
A
OFF
ON
OFF
t = 0
1
2
3
4 seconds
visual stimulus
full field flash
M1
M5
M9
M10
average projection ON
(t = 1-1.6 seconds)
average projection OFF
(t = 3-3.6 seconds)
overlay ON OFF
B
full field flash responses
ROIs (axons terminals, M9/10)
1 2 3 4 5 6 7
ROI
ΔF/F
seconds
C
receptive field mapping
bar direction
ROI
bar position (y)
visual stimulus screen
RF centers
M1
M5
M9/10
visual stimulus
moving bar
bar direction
bar position (x)
D
average full field flash responses
binned by RFC position
on-screen RFCs
("normal")
off-screen RFCs
("inverted")
compartment
M1
M5
M9/10
seconds
response amplitude
M1 M5 M10
compartment

**Figure S3: Mi1 receptive field mapping.** A: Average projections of jRGECO1a signals in Mi1 neurons while the screen was light (ON, t = 1-1.6s, left) or dark (OFF, t = 3-3.6s, middle); image on the right shows the overlay of the ON (green) and OFF (magenta) images. B: Cropped image corresponding to the dashed white box on the overlay image in A; average responses corresponding to the ROIs indicated on the images are plotted over time. Note that light increments drove cytosolic calcium increases in ROIs 1-4 and calcium decreases in ROIs 5-7. C: Line plots show jRGECO1a responses to white bars moving either vertically (top left) or horizontally (bottom right) in ROI 2 and 7 from the image in B. For ROIs that responded to the moving light bars, peak jRGECO1a signals were used to map receptive field centers (RFCs) on the stimulus screen (top right); RFCs for ROIs 1-4 are shown shades of blue and green. D: Average full field flash responses for ROIs with RFCs mapped to the stimulus screen ("on-screen RFCs") and ROIs that did not map to the stimulus screen ("off-screen RFCs"). The full field flash stimulus drove cytosolic calcium increases in on-screen ROIs ("normal" responses for Mi1, which are ON neurons) and calcium decreases in off-screen ROIs ("inverted" responses, consistent with activation of an inhibitory surround).

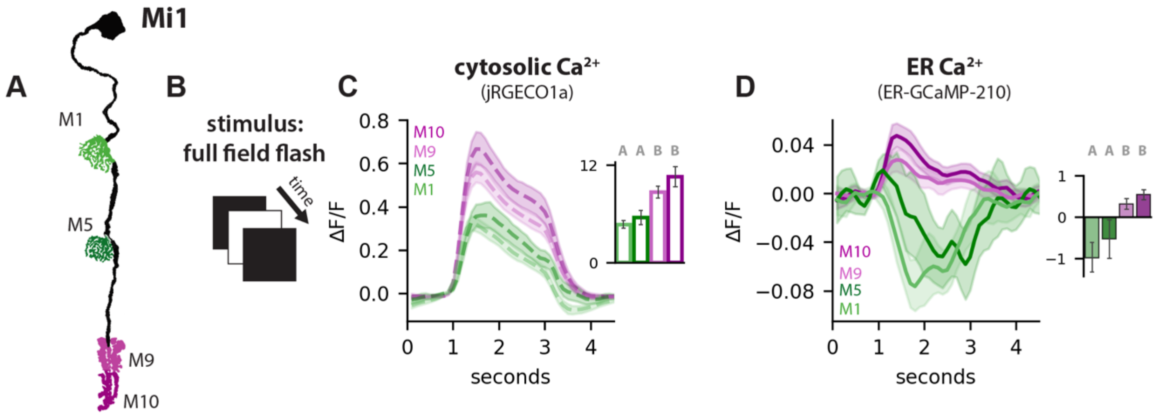


**Figure S4: ER and cytosolic calcium responses in all layers of Mi1.** A: Mi1 neurons, with dendrite compartments (M1 and M5) in shades of green and axon compartments (M9 and M10) in shades of magenta. B: Full field flash stimulus. C-D: Average cytosolic calcium (C) and ER calcium (D) responses to full field flash stimuli. Insets show average response amplitudes; N = 101 ROIs from 8 flies (M1), 60 ROIs from 8 flies (M5), 244 ROIs from 13 flies (M9), 148 ROIs from 13 flies (M10). Letters above bar plots indicate groups that are significantly different (Kruskal Wallis with post-hoc Dunn's test, $p < 0.05$).

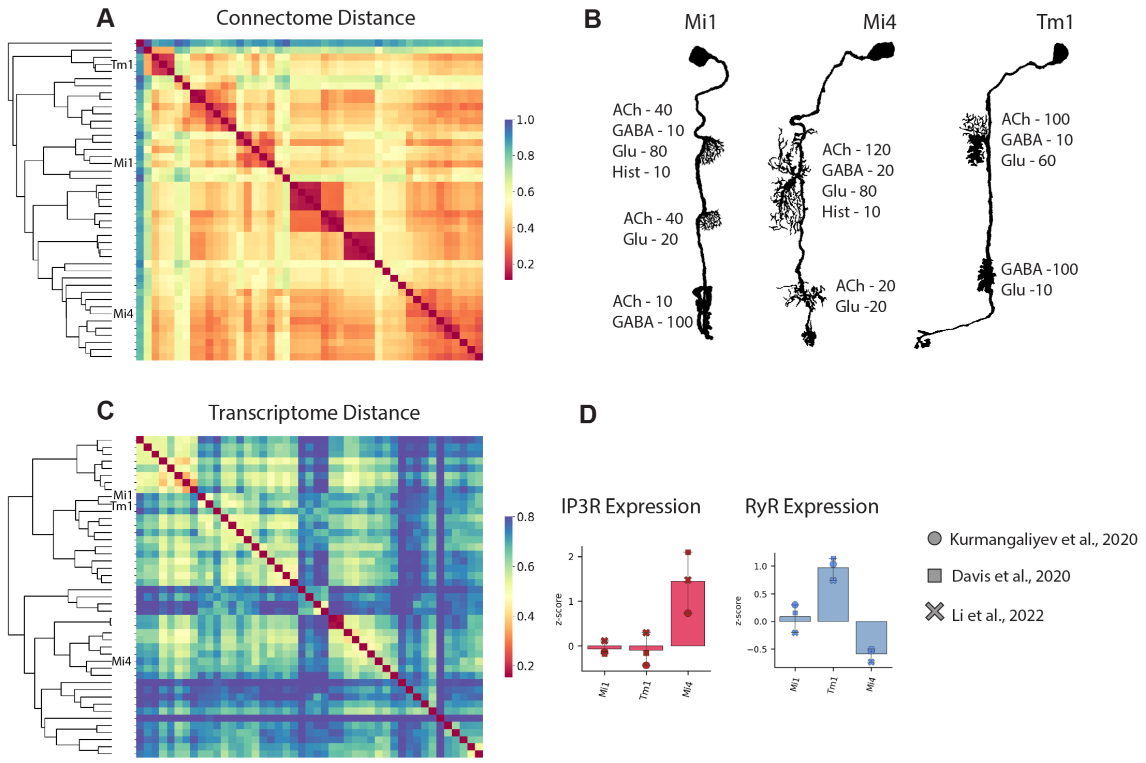


**Figure S5: Mi1, Mi4, and Tm9 have diverse morphologies, synaptic inputs, and transcriptomes.** A: *Drosophila* medulla neurons clustered based on synaptic inputs[1]; note that Mi1, Mi4, and Tm1 occupy distinct clusters. B: Average number of cholinergic, GABAergic, glutamatergic, and histaminergic inputs onto each subcellular compartment in Mi1, Mi4, and Tm1 neurons[1]. Synapse numbers rounded to the nearest 10. C: *Drosophila* medulla neurons clustered based on ssRNAseq data[2]. D: Average normalized expression of IP3R and RyR in Mi1, Mi4, and Tm1; symbols indicate expression levels in three different datasets[2-4].
